# Therapeutic doses of ketamine acutely attenuate the aversive effect of losses during decision-making

**DOI:** 10.1101/2022.07.25.501468

**Authors:** Mariann Oemisch, Hyojung Seo

## Abstract

The discovery of rapid-acting antidepressant, ketamine has opened a pathway to a new generation of treatments for depression, and inspired neuroscientific investigation based on a new perspective that non-adaptive changes in the intrinsic excitatory and inhibitory circuitry might underlie the pathophysiology of depression. Nevertheless, it still remains largely unknown how the hypothesized molecular and synaptic levels of changes in the circuitry might mediate behavioral and neuropsychological changes underlying depression, and how ketamine might restore adaptive behavior. Here, we used computational models to analyze behavioral changes induced by therapeutic doses of ketamine, while rhesus macaques were iteratively making decisions based on gains and losses of tokens. When administered intramuscularly or intranasally, ketamine reduced the aversiveness of undesirable outcomes such as losses of tokens without significantly affecting the evaluation of gains, behavioral perseveration, motivation and other cognitive aspects of learning such as temporal credit assignment and time scales of choice and outcome memory. Ketamine’s potentially-antidepressant effect was separable from other side effects such as fixation errors, which unlike outcome evaluation, was readily countered with strong motivation to avoid errors. We discuss how the acute effect of ketamine to reduce the initial impact of negative events could potentially mediate longer-term antidepressant effects through mitigating the cumulative effect of those events produced by slowly decaying memory, and how the disruption-resistant affective memory might pose challenges in treating depression. Our study also invites future investigations on ketamine’s antidepressant action over diverse mood states and with affective events exerting their impacts at diverse time scales.

## Introduction

Depression is a debilitating mental illness that causes enormous personal sufferings as well as societal and economic burden (1-3). The discovery of ketamine, a non-competitive *N*-methyl-*D*-aspartate (NMDA) receptor antagonist as a rapid-acting antidepressant, inspired extensive research to understand the mechanisms of ketamine’s action as well as a new perspective on the neurobiology of depression that depression might reflect non-adaptive changes in the glutamatergic and GABA (Gamma Aminobutyric Acid) -ergic signal transmission of the prefrontal and limbic system in response to environmental stress and adversity (4-13). Despite recent progress in understanding the mechanisms of ketamine’s antidepressant action at cellular, synaptic and network levels, many questions remain regarding enhancing the specificity to target depressive symptoms without undesirable side effects as well as long-term safety for repeated dosing (14-17).

A major gap in our knowledge that hampers the effort to design novel treatments with high specificity and low off-target effect lies at the functional-systems level to link cellular and synaptic phenomena to physiological and behavioral processes that directly cause the symptoms of depression. In light of symptomatic heterogeneity within the diagnostic category of mental disorders including depression, the importance of integrative and comprehensive analyses with regard to multiple functional domains/systems has been recognized to promote interpretable and translatable results from pre-clinical and clinical research (18, 19). Nevertheless, this type of analyses and results for assessing the behavioral effect of ketamine’s antidepressant action have seldom been available from the behavioral tests prevalently used in rodents (e.g. forced swim test, sucrose preference test etc.) or self-reports from human subjects/patients. Another type of scarce data is early behavioral signs of ketamine’s action that might mediate and/or predict its longer-term effects. Ketamine has shown to induce rapid-onset (<1 hour) cellular, molecular, physiological and connectivity changes that are thought to reflect and/or mediate its antidepressant action (20-25). By contrast, the acute behavioral effect that can be indicative of and/or predict longer-term effects has been hard to isolate from dissociative side effects.

Here, we assessed the effects of therapeutic doses of ketamine (0.5-1mg/kg) administered intramuscularly or intranasally on cognitive, affective and motivational aspects of learning and decision making. Three rhesus monkeys performed a token-based biased matching pennies game against a computerized opponent in which they gained or lost tokens as the outcome of their decisions (26, 27). We found that ketamine acutely reduced the adverse effect of undesirable outcomes such as losses of tokens, which was independent of other side effects such as ocular nystagmus. However, ketamine did not significantly change animals’ motivation to earn reward, behavioral perseveration or other cognitive aspects of learning and memory. These results constitute a rare report of an acute effect of therapeutic dose of ketamine on the processing of affectively negative events during dynamic decision-making. Our study also demonstrates the sensitivity and the potential of the non-human primate model in assessing behavioral and systems-level mechanisms of antidepressant action as well as investigating the pathophysiology of depression when combined with recently developed tools for large-scale neural recording and circuit manipulation.

## Methods and Materials

### Animal preparation

Three male monkeys (Macaca mulatta, P, Y and B; body weight, 12-16 kg) were used. The animal was seated in a primate chair with its head fixed, and eye position was monitored with a high-speed eye tracker (ET 49, Thomas Recording, Giessen, Germany). All procedures used in this study were approved by the Institutional Animal Care and Use Committee at Yale University, and conformed to the Public Health Service *Policy on Humane Care and Use of Laboratory Animals* and *Guide for the Care and Use of Laboratory Animals*.

### Behavioral task

Animals were trained to perform a token-based biased matching-pennies (BMP) task (Figure 1, A). Details of the task have been previously published (26). Briefly, animals played a competitive BMP game against a computerized opponent and the outcome of each trial was determined jointly by the animal’s and the opponent’s choice according to the payoff matrix (Figure 1, B). The payoff matrix was adjusted for each animal to produce comparable choice behavior across all three animals. When the animals matched the opponent’s choice, they gained one (monkey P) or two (monkey Y/B) tokens, whereas non-matching choices incurred either zero token (i.e. neutral outcome) or loss of one (Y/B) or two (P) tokens, with the probability of loss being higher (lower) than that of zero token for the risky (safe) target. The locations of “safe” and “risky” targets were fixed during a block of 40 trials and then switched with a probability of 0.1 afterwards. The computer opponent simulated a rational player who exploits any predictable patterns in the animal’s choice and outcome history to minimize the animal’s payoff (26). To maximize payoff, the animals were required to dynamically change their choices to minimize serial correlations in the sequence of choice and outcome that could be exploited by the opponent.

**Figure 1.**
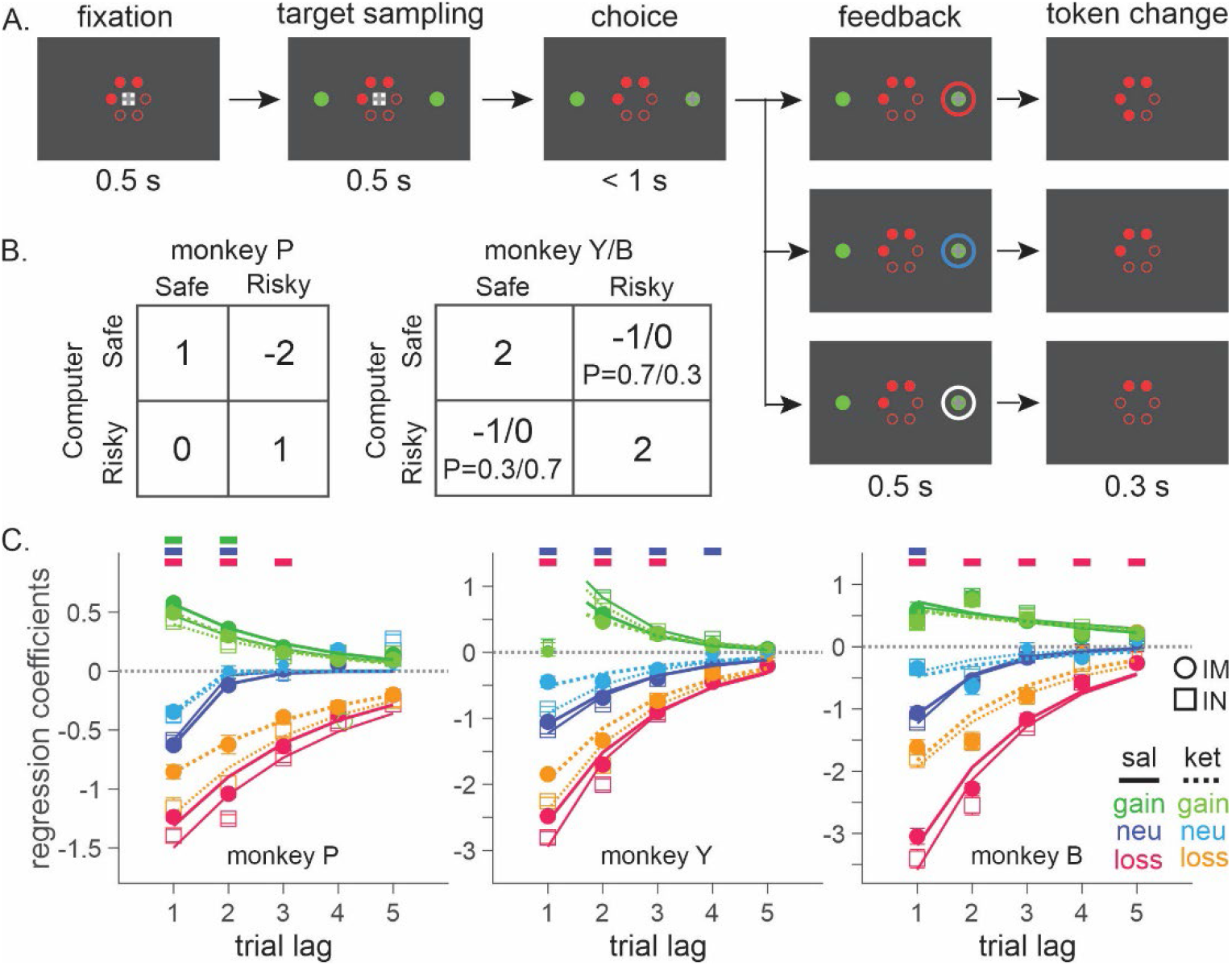
Biased matching pennies (BMP) task and ketamine-induced behavioral modulation. **A**. Temporal sequence of trial events. Gray cross indicates the target that the animal was required to fixate during each epoch. Each trial began with the animal’s gaze on the fixation target displayed on the center of a computer monitor for 0.5-s. Solid red disks around the fixation target indicated the tokens owned by the animal, namely asset, with empty disks serving as placeholders for the tokens to be acquired for exchange with juice reward. After two green disks were displayed for 0.5-s at the diametrically opposed positions along the horizontal meridian, the fixation target was extinguished signaling the animal to indicate its choice by shifting gaze to one of the two green disks. After 0.5-s of fixation, a feedback ring appeared around the chosen target with its color indicating the choice outcome, followed by the corresponding change in tokens. Once the animal collected 6 tokens, they were automatically exchanged with 6 drops of apple juice. After juice reward, the animal began the subsequent trial with 2-4 free tokens. Tokens and placeholders stayed on the screen throughout the trial and inter-trial interval. **B**. Payoff matrix of BMP. **C**. Coefficients from logistic regression models applied separately to the data from saline (sal) and ketamine (ket) sessions with intramuscular (IM) and intranasal (IN) administration. Lines are exponential functions best fit to the coefficients from saline (solid) and ketamine (dotted) sessions, and those from IM (thick) and IN (thin) sessions. Bars at the top of each panel indicate that the difference between saline and ketamine sessions is statistically significant with the horizontal position and color of each bar coding trial lag and outcome (same color scheme as that of saline sessions), respectively.

### Administration of ketamine

For intramuscular (IM) administration, 0.25mg/kg or 0.5mg/kg of ketamine (Ketaset®, Zoetis Inc.) was injected into the right or left calf muscle. Doses were selected based on the therapeutic doses of ketamine when it is used as antidepressant for human patients. For intranasal (IN) administration, 0.5mg/kg or 1mg/kg of ketamine was delivered into the right or left nostril using an intranasal mucosal atomization device (MAD Nasal™, Teleflex Medical). Ketamine was administered with a 6.4-day interval on average (2-7 days), and sterile saline (IM) or sterile water (IN) were administered in-between successive ketamine sessions.

### Analysis of choice behavior

#### Logistic regression model

The effect of past outcomes on an animal’s choice was analyzed using the following model:

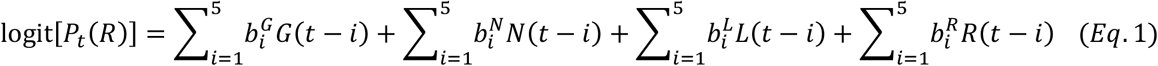

where *P*_*t*_ (*R*) is the probability of animal’s choosing right target at the current trial *t, G(t-i), N(t-i), L(t-i) and R(t-i)* are regressors encoding gain, neutral, loss and juice reward outcome that occurred after right (left) choice in the *i*-th trial into the past as 1(-1), and other outcomes as 0, respectively. 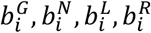 are regression coefficients, with positive (negative) coefficients indicating that the particular outcome at trial t-*i* tended to increase (decrease) the animal’s tendency to choose the same target at trial *t*, relative to chance level (Figure 1, C). Separate regression models were fit to saline, intramuscularly (IM)- and intranasally (IN)-administered ketamine sessions for each animal.

#### Nonlinear regression - exponential model

We modeled the temporal decay of the effect of past outcome over multiple trials and its modulation by ketamine with the following exponential function:

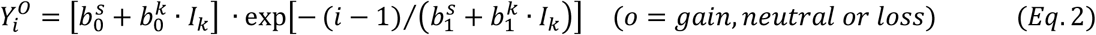

where *i* is trial lag (*i*-th trial into the past), *I*_*k*_ is an indicator variable encoding ketamine (saline) session as 1 (0). 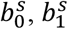 are initial amplitude and decay time constant respectively, for saline session, whereas 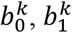 are modulation by ketamine. Exponential functions were fit to the data with nonlinear regression using *fitnlm*.*m* in *MATLAB* (MathWorks), separately for each outcome.

#### Reinforcement learning models

We used reinforcement learning (RL) models to investigate the potential cognitive/computational processes that might be modulated by ketamine. RL models have been extensively and successfully applied to explain reward-guided choice behavior (28). The general idea of RL is that the outcomes experienced from a particular action/state are integrated over time to form the reward expectation or the value of that action/state, and that an action among alternatives is chosen to maximize the expected outcome. One algorithm for the integration is to incrementally update the old expectation/value with a prediction error which can be formalized as follows for a single-state situation (29):

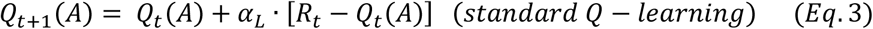

where *Q*_*t*_ (*A*) is the value function for action A at trial *t, α*_*L*_ is learning rate and *R*_*t*_ − *Q*_*t*_ (*A*) is the discrepancy between actual outcome,*R*_*t*_ and the expectation/value, namely reward prediction error. This formula can be rearranged as follows (30, 31):

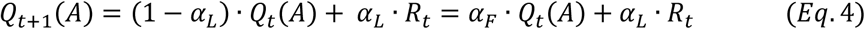

where *α*_*F*_ and *α*_*L*_ are separate parameters for the forgetting rate of the old value estimate and the learning rate of a new outcome, respectively. With 0 < *α*_*F*_ < 1, the old value function decays exponentially over time, simulating forgetting. Action selection was modeled with a softmax function,

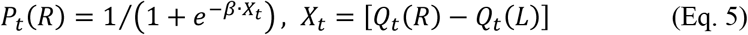

where probability of choosing right target, *P*_*t*_ (*R*) was a logistic function of the estimated value of right relative to left choice. Inverse temperature, *β* regulated randomness of choice determining the influence of action values on decision.

To investigate the effects of ketamine, we first tested multiple variants of the general model and determined the model that best explained normal behavior during saline sessions (Table 1; Figure 2). This is important, since we can expect to characterize the effect of ketamine accurately only in the context of a reliable behavioral model for how the animal’s behavior was influenced by the outcomes of their previous choices. Then, we examined how the parameters of this model were modulated by ketamine (Table 2; Figure 3).

**Table 1.** Reinforcement learning models for normal behavior during saline sessions.

| model | value update |
| --- | --- |
| **Q-learning (Q)** | $Q_{t+1}(A) = \alpha_F \cdot Q_t(A) + (1 - \alpha_F) \cdot R_t$, if A is chosen.<br>$R_t = 1, 0, -1$ for gain, neutral and loss, respectively |
| **Q-learning with subjective outcome evaluation (QSE)** | $Q_{t+1}(A) = \alpha_F \cdot Q_t(A) + \Delta_t$, if A is chosen.<br>$\Delta_t = \Delta_G, \Delta_N, \Delta_L$ for gain, neutral and loss (QSE)<br>$\Delta_N=0$ (QSE-R) |
| **Differential forgetting (DF)** | $Q_{t+1}(A) = \alpha_{F-C} \cdot Q_t(A) + \Delta_t$, if A is chosen,<br>$Q_{t+1}(A) = \alpha_{F-UC} \cdot Q_t(A)$, if A is not chosen.<br>$\Delta_t = \Delta_G, \Delta_N, \Delta_L$ for gain, neutral and loss (DF)<br>$\Delta_N=0$ (DF-R) |
| **Non-differential forgetting (NDF)** | $Q_{t+1}(A) = \alpha_F \cdot Q_t(A) + \Delta_t$, if A is chosen,<br>$Q_{t+1}(A) = \alpha_F \cdot Q_t(A)$, if A is not chosen.<br>$\Delta_t = \Delta_G, \Delta_N, \Delta_L$ for gain, neutral and loss (NDF)<br>$\Delta_N=0$ (NDF-R) |
| **DF with asset-gated outcome evaluation (DF-A)** | $Q_{t+1}(A) = \alpha_{F-C} \cdot Q_t(A) + \lambda \cdot asset_t \cdot R_t$, if A is chosen,<br>$Q_{t+1}(A) = \alpha_{F-UC} \cdot Q_t(A)$, if A is not chosen.<br>$R_t = 1, 0, -1$ for gain, neutral and loss, respectively |

**Table 2.**
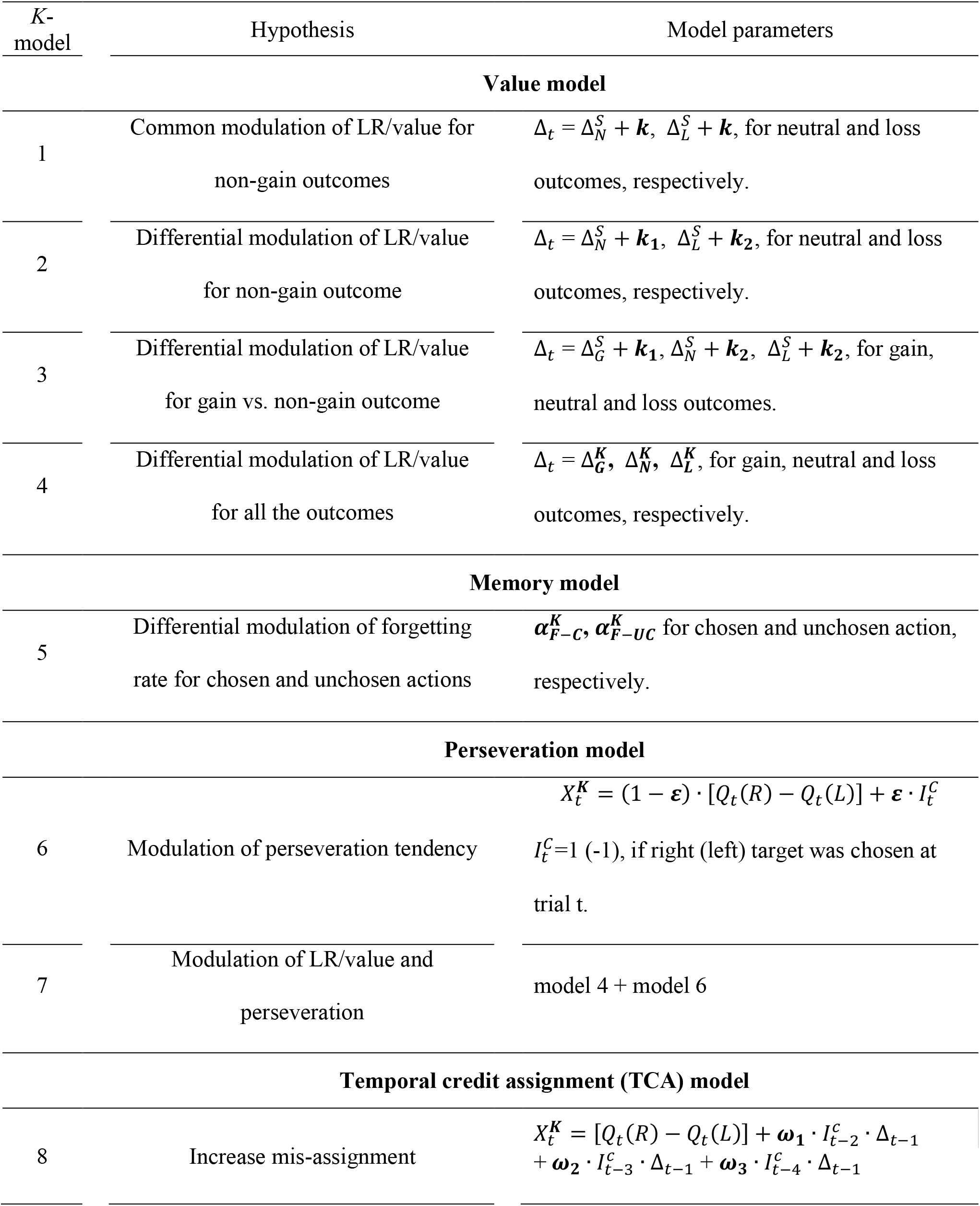
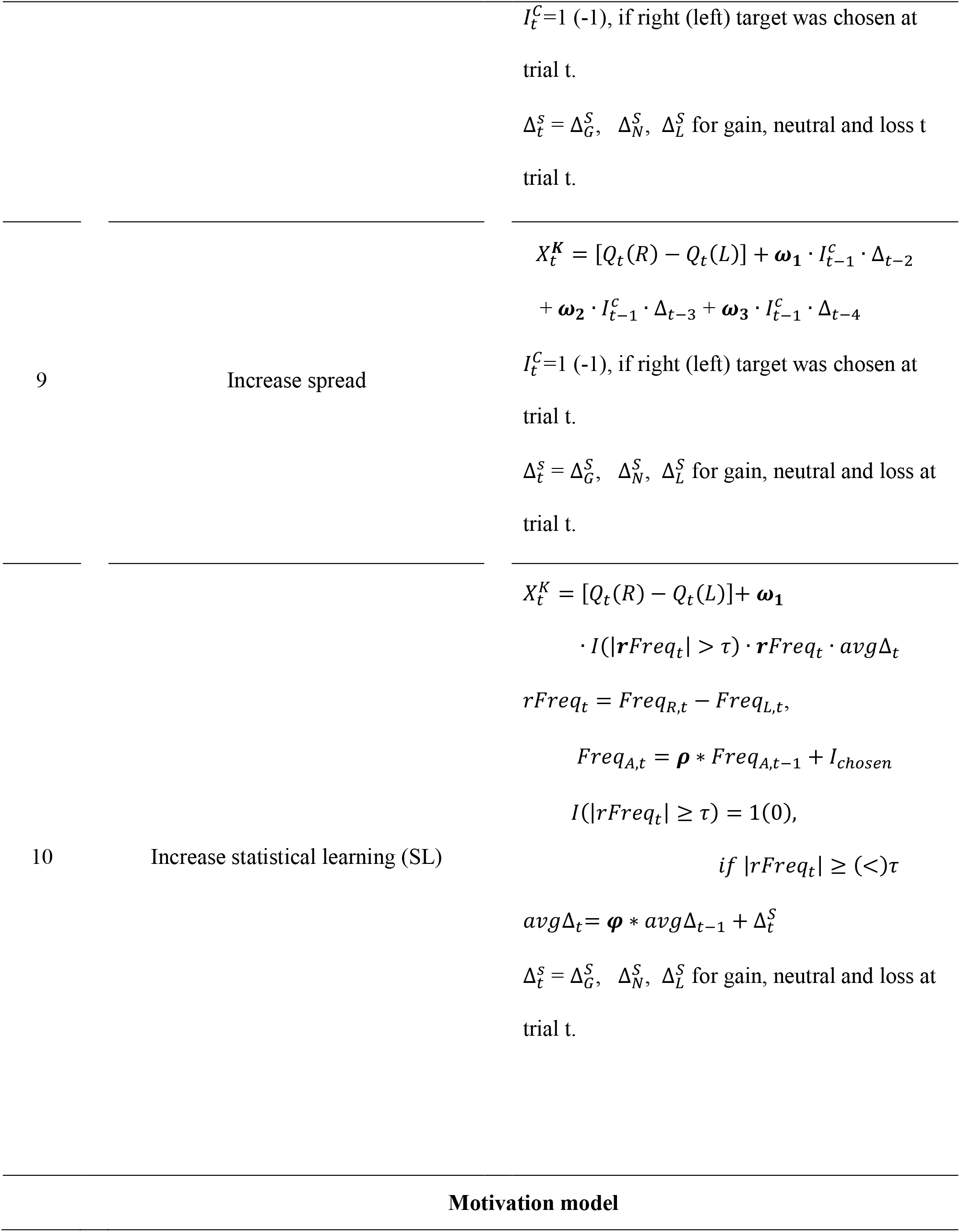
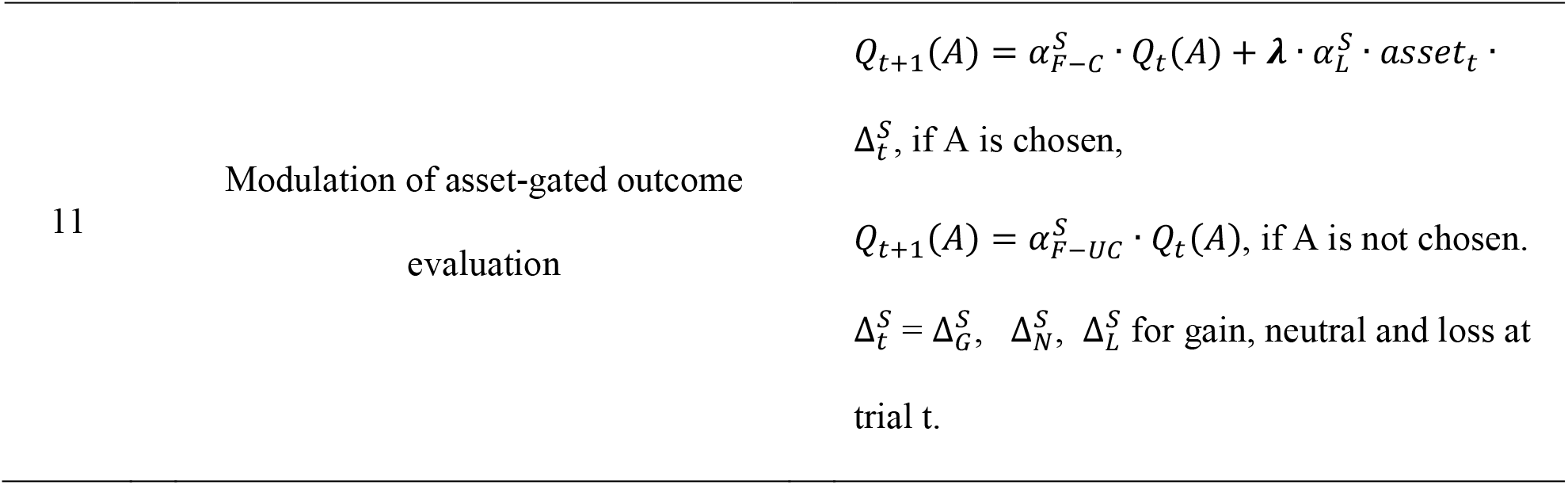
Reinforcement learning models for ketamine-induced modulation of choice behavior.

**Figure 2.**
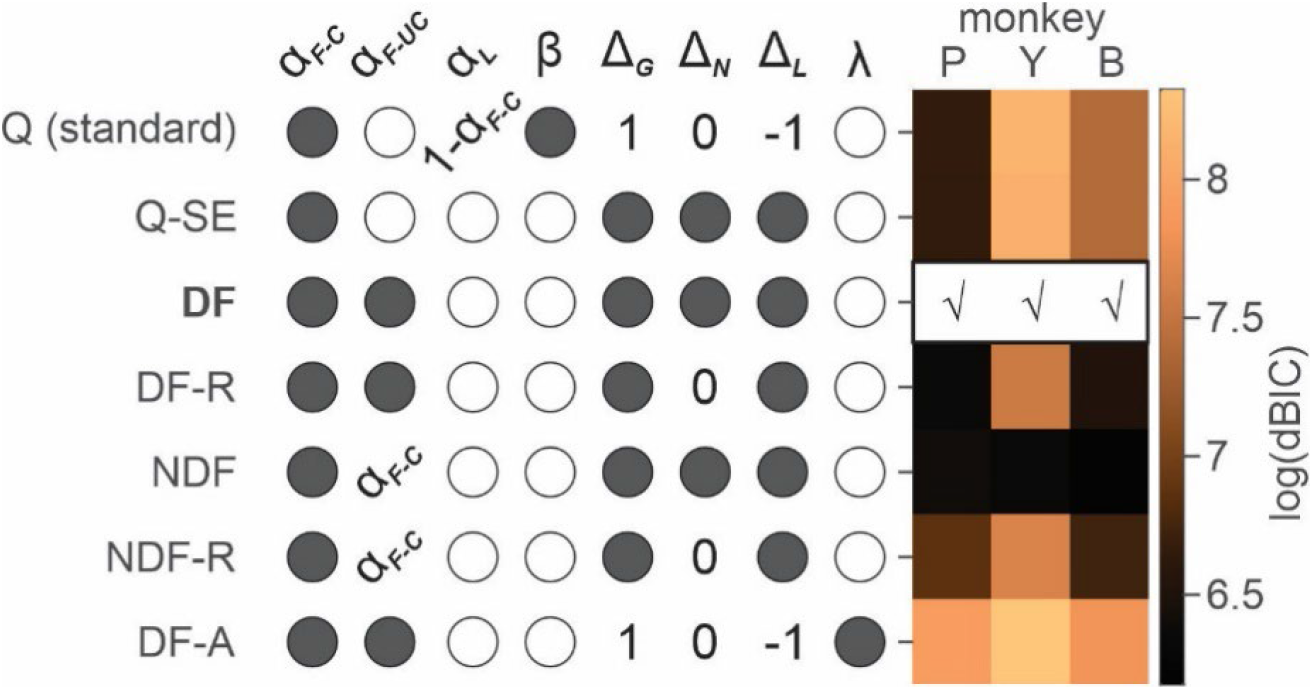
Model comparison for saline sessions. Each row shows free (filled circles), unused (empty circles) and fixed (specific values) parameters of a variant of reinforcement learning (RL) model. For the details of RL models and parameters, see Table 1. Heat map on the right represents natural logarithm of differential BIC (Bayesian Information Criterion) of each variant from the best model. Best model for each animal is indicated by check (√) mark. *Q(standard)*: standard Q-learning model; *Q-SE*: Q-learning with subjective outcome evaluation; *DF*: differential forgetting model; *DF-R*: differential forgetting with neutral outcome being the reference point; *NDF*: non-differential forgetting model; *NDF-R*: non-differential forgetting model with neutral outcome being the reference point; *NDF-A*: non-differential forgetting model with asset-gated outcome evaluation.

**Figure 3.**
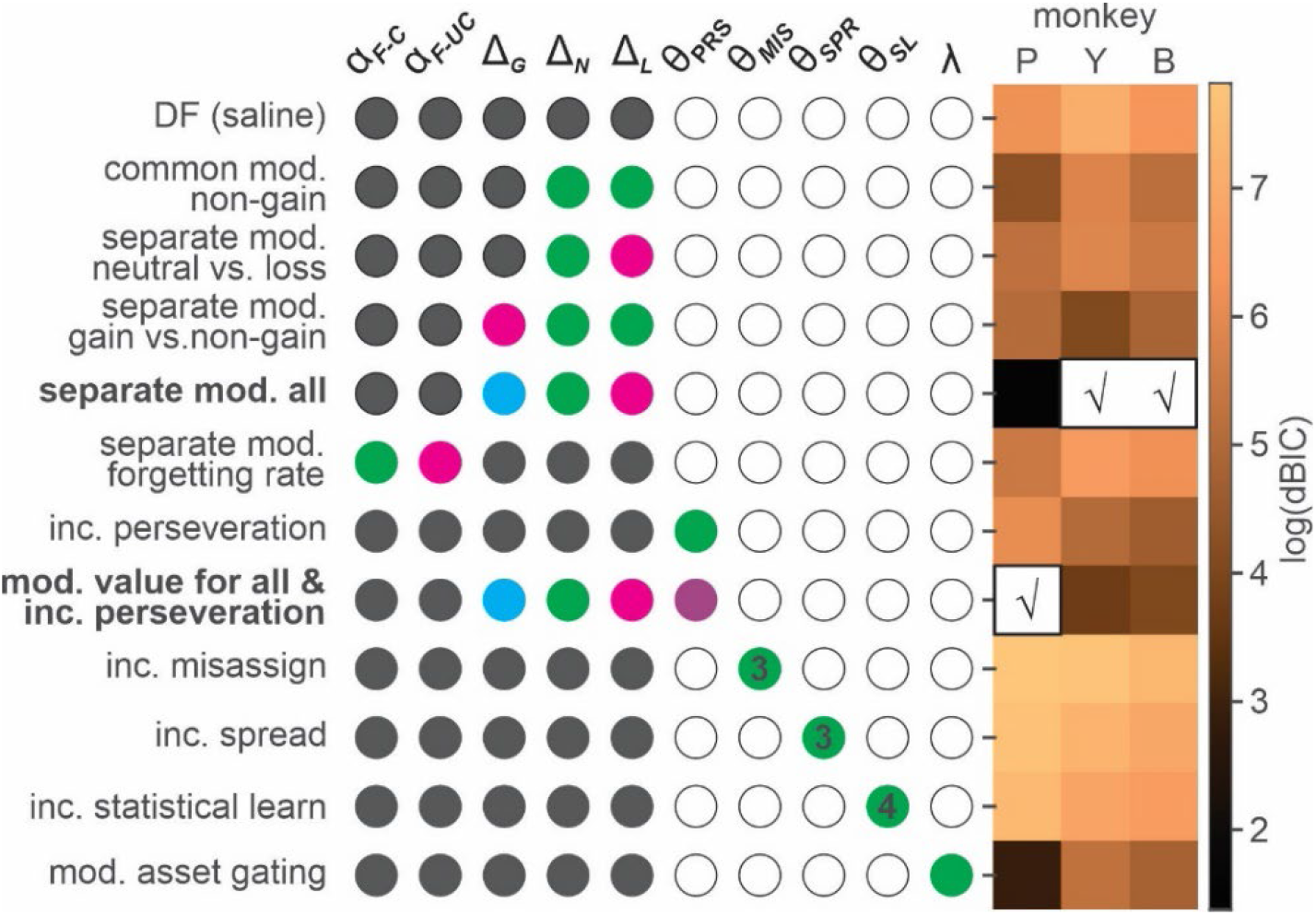
Model comparison for the effects of ketamine. Each row shows fixed (gray-filled circles), free (colored circles) and unused (empty circles) parameters of a variant of differential forgetting (DF) model. For the details of each model, see Table 2. θ represents a set of parameters added to a particular variant of DF model, and the number inside colored circles indicate the number of added parameters (>1). For the details of each model, see Table 2. Heat map on the right represents natural logarithm of differential BIC (Bayesian Information Criterion) of each variant from the best model. Best model for each animal is indicated by check (√) mark. *DF(saline)*: differential forgetting model fit to the data from saline session; *Common mod. non-gain*: common modulation of non-gain outcome evaluation; *Separate mod. neutral vs. loss*: differential modulation of neutral and loss outcome evaluation; *Separate mod. gain vs. non-gain*: differential modulation of gain and non-gain outcome evaluation; *Separate mod. all*: outcome-dependent modulation of outcome evaluation; *Separate mod. forgetting rate*: differential modulation of forgetting rate for chosen and unchosen target, *Inc. perseveration*: increased perseveration; *Mod. value for all & inc. perseveration*: outcome-dependent modulation of outcome evaluation and increased perseveration; *Inc. misassign*: increased backward credit misassignment; *Inc. spread*: increased forward spread in credit assignment; *Inc. statistical learning*: increased statistical learning of reward rate; *Mod. asset gating*: modulation of asset-gated outcome evaluation.

#### Model selection for saline sessions

In the most restricted, standard Q-learning model, value was updated only for the chosen action by taking a weighted average of *Q*_*t*_ (*A*) and *R*_*t*_ with *α*_*F*_ = 1 − *α*_*L*_. *R*_*t*_, was fixed as -1, 0 and 1 for gain, neutral and loss outcome, respectively. These arrangements confined action value Q within the range of -1 to 1.

For three variants of RL models - Q-learning with subjective evaluation (QSE), differential forgetting (DF) and non-differential forgetting (NDF) models, we relaxed the restrictions of the Q-learning model in two ways. *First, R*_*t*_ was set as a free parameter reflecting subjective outcome evaluation. When both *α*_*L*_ and *R*_*t*_ are set as free parameters, the model becomes underdetermined. Therefore, in these models, the product of learning rate and outcome value, *α*_*L*_ · *R*_*t*_ was consolidated into a single parameter Δ_*t*_ or the total amount of change due to a new outcome which could take different values for gain (Δ_*G*_), neutral (Δ_*N*_) and loss (Δ_*L*_) outcome. We also tested the models in which Δ_*N*_ was fixed as 0 simulating effectively neutral outcome. *Second*, action value Q for the unchosen action was allowed to decay, with differential forgetting rates for chosen (*α*_*F*−*C*_ ) and unchosen (*α*_*F*−*UC*_ ) actions in a DF model, and with a common forgetting rate, *α*_*F*_ in an NDF model. Action values were updated only for the chosen action in a QSE model.

Finally, we tested an additional model in which outcome evaluation was gated by asset (non-differential forgetting with asset-gated outcome evaluation - NDF-A model) or the number of tokens owned by the animal at the time of decision, simulating temporal discounting of outcome value depending on its temporal distance to the primary reward that was always given only after 6 tokens were acquired.

For all tested models other than Q-learning model, *β* was fixed to be 1, as a model was underdetermined when both *β* and Δ_*t*_ could vary freely. Model parameters were fit to choice data to maximize the likelihood of data. Bayesian information criterion (BIC) was used for model comparison and selection of the best model.

#### Model selection for Ketamine sessions

To understand the effect of ketamine, we started with the best-fitting model of normal behavior during saline sessions – the differential forgetting (DF) model, and then asked how ketamine might modulate the parameters of this model. The null (baseline) model assumed that ketamine didn’t cause any parametric changes in the behavior, and the model prediction of trial-to-trial choice was generated with the maximum likelihood (ML) parameters, namely “baseline parameters” estimated from saline sessions as follows:

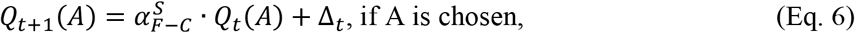

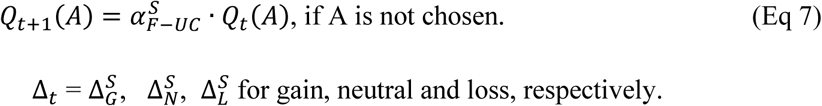

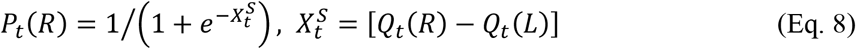

where 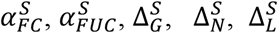 (baseline parameter-estimates) were ML parameters of the DF model fit to the data from saline sessions.

In each of the ketamine models, we set a subset of parameters of the DF model to freely vary from the baseline parameter-estimates. *Value models* (*K*-model 1-4, Table 2) hypothesized that ketamine changed outcome evaluation differentially for gain, neutral and loss outcomes, whereas a *memory model* (*K*-model 5) tested whether ketamine affected the forgetting rate or the rate of value decay. *Perseveration models* hypothesized that ketamine affected the behavioral tendency to perseverate (*K*-model 6), possibly in tandem with a change in value update (*K*-model 7). *Temporal credit assignment (TCA) models* tested whether ketamine changed the way a particular outcome properly updated the value of causative action, possibly leading to misassignment of the credit for an outcome to the choice that happened in the past (*K*-model 8), spread of the credit to future choice (*K*-model 9) or assignment of the credit for recent outcomes collectively to the frequently chosen target in the near past (*K*-model 10) (30, 31). Finally, a *motivation model* (*K*-model 11) hypothesized that ketamine changed the gain with which asset position modulated the value update for each outcome.

### Analysis of ketamine-induced ocular nystagmus

Ocular nystagmus during fixation on a peripheral target consists of two components - slow centripetal drift (i.e. slow phase), and a subsequent corrective saccade to re-capture the target (i.e. fast phase) (34). Eye position was sampled and recorded at 250Hz. We used average velocity of eye movement during pre-feedback fixation on a chosen target to detect the slow phase of the nystagmus induced by ketamine. The sign of eye velocity was adjusted in the way that centripetal eye movement during fixation had negative velocity. Fast phase eye movement with velocity >20 ° (visual angle)/s was first removed from the pre-feedback fixation period, and then eye velocity was averaged during the remaining time points of the fixation period for each trial (for statistical analysis) or in 1-min non-overlapping bins for visualization of time course (Figure 4).

**Figure 4.**
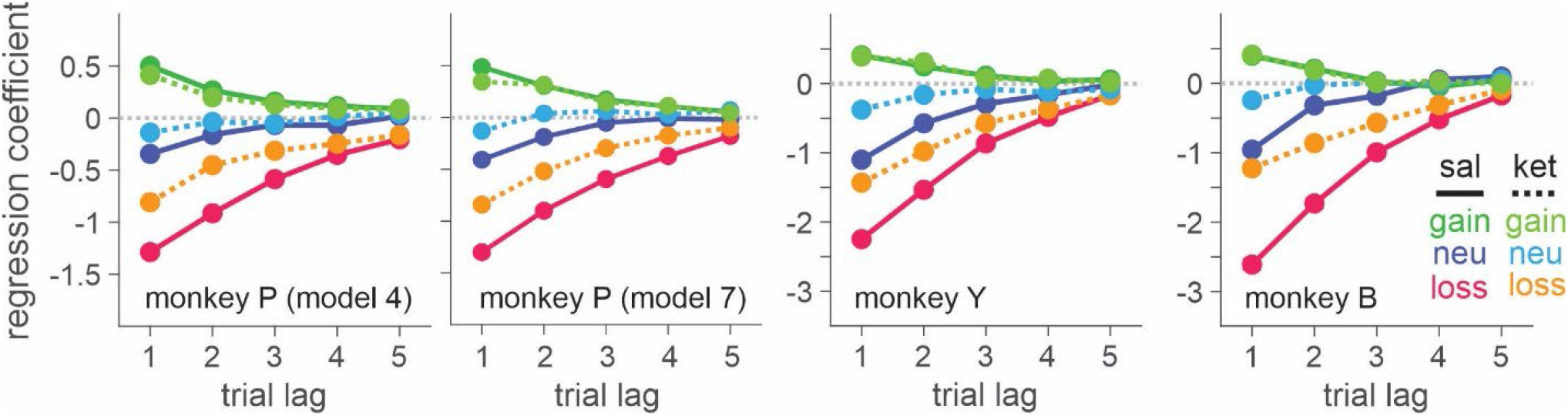
Ketamine-induced behavioral modulation simulated with differential forgetting model (for saline session) and best-fitting *K*-model (for ketamine session). Simulated data was generated with the maximum likelihood parameters of best-fitting models for saline (differential forgetting model) and ketamine (*K*-model 4: differential modulation of value for all the outcomes) sessions. Simulated choice was analyzed with logistic regression model (Eq. 1).

### Results

Three rhesus monkeys were trained to play a token-based matching pennies (BMP) game against a computerized opponent, in which their choice typically depended on the outcomes obtained from each choice in the past trials (Figure 1, A, B). In the control sessions where saline was administered, animals tended to repeat the choice that yielded a token, while switching away from the choice that led to non-gain outcomes, with a stronger tendency to switch after loss than neutral outcome (i.e. zero token) (Eq. 1; Figure 1, C). The reinforcing and punishing effects of gain and non-gain outcomes also decayed exponentially over time, with the recent outcomes exerting a larger effect than remote ones. Ketamine selectively reduced the punishing effect of non-gain outcomes, with significantly larger change being induced after loss than neutral outcome (Figure 1, C). In addition, the ketamine-induced modulation was larger for the effect of recent than remote outcomes (*ANOVA*, 3-way interaction among outcome, trial-lag and drug condition, *F(8, 1964)*=4.05, *F(8, 1649)*=4.46, *F(8, 509)*=3.62 for animal P, Y, B, respectively. *p*<.001 in all animals for intramuscular (IM) administration of 0.5mg/kg; *F(8, 1229)*=4.89, *p*<.001 in animal P; *F(8, 389)*=1.89, *p*=.06 in Y; *F(8, 1002)*=3.27, *p*=.001 in B for intranasal (IN) administration of 1mg/kg). 0.25mg/kg of IM-administered, and 0.5mg/kg of IN-administered ketamine produced qualitatively similar results (Supplemental Figure S1). We focused our main analyses on the data collected with 0.5mg/kg IM- and 1mg/kg IN-administered ketamine, which produced comparable plasma concentration and behavioral effects (Supplemental Information and Figure S2). Plasma concentration and its time course over 60 minutes were also comparable to those measured after 0.5mg/kg in human subjects (35).

Outcome-dependent choice behavior and its modulation by ketamine were also parametrically modeled by exponential function, in which the initial impact of each outcome at the time of delivery gradually decayed over subsequent trials (Eq 2; Figure 1, C). Ketamine significantly reduced initial amplitude/impact of non-gain outcomes 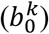 consistently in all three animals, and with both IM 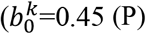, 0.63 (Y), 1.36 (B) for loss, *t*-test, *p*<10^−11^ for all animals; 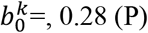, 0.58 (Y), 0.61 (B) for neutral outcome, *p*<10^−5^ for all animals) and IN administration 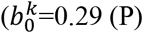, 0.56 (Y), 1.36 (B) for loss, *p*<10^−11^ for all animals; 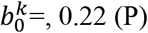, 0.31 (Y), 0.83 (B) for neutral outcome, *p*<.001 for all animals). Ketamine didn’t significantly modulate the time constant of decay, and the modulation of initial impact was sufficient to explain the behavioral phenomena.

Dependence of choice on the cumulative effect of temporally discounted past outcomes is the signature of reinforcement learning (RL) (28). We used RL models to gain further insights into the possible mechanisms underlying ketamine-induced behavioral modulation (see Methods and Materials). We first determined the variant/class of RL model that best explained the animals’ choice behavior under normal conditions (i.e., saline sessions) (Table 1). The differential forgetting (DF) model best fit the data in all three animals, in which the value functions of chosen and unchosen action decayed at different rates, with forgetting being slower for chosen than unchosen action (Figure 2, Table 3).

**Table 3.**
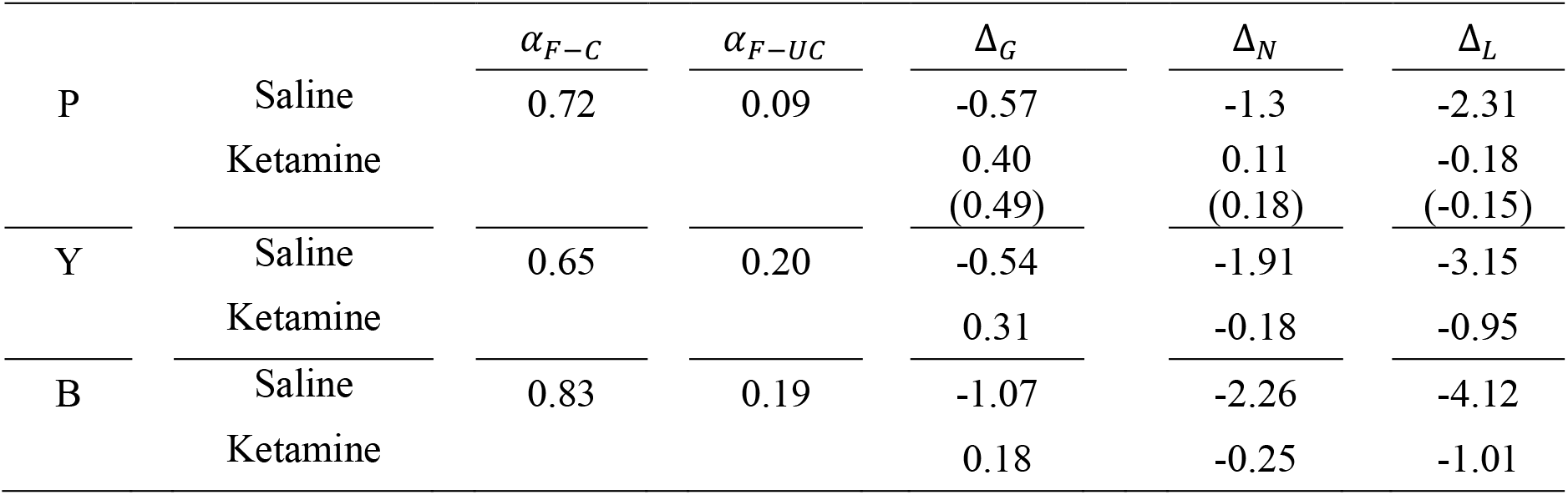
Maximum likelihood parameter estimates of the best models for saline and ketamine sessions.

| | | $\alpha_{F-C}$ | $\alpha_{F-UC}$ | $\Delta_G$ | $\Delta_N$ | $\Delta_L$ |
| --- | --- | --- | --- | --- | --- | --- |
| P | Saline | 0.72 | 0.09 | -0.57 | -1.3 | -2.31 |
|  | Ketamine |  |  | 0.40<br>(0.49) | 0.11<br>(0.18) | -0.18<br>(-0.15) |
| Y | Saline | 0.65 | 0.20 | -0.54 | -1.91 | -3.15 |
|  | Ketamine |  |  | 0.31 | -0.18 | -0.95 |
| B | Saline | 0.83 | 0.19 | -1.07 | -2.26 | -4.12 |
|  | Ketamine |  |  | 0.18 | -0.25 | -1.01 |

Building on the best model explaining normal behavior – the DF model, we asked how ketamine (0.5mg/kg, IM) modulated the parameters of this model by testing variants of DF models, in each of which a different subset of parameters of the model were allowed to vary/deviate freely from the baseline parameters estimated from saline sessions (Table 2).

In all three animals, the model incorporating valence-dependent change in outcome evaluation best fit the choice data from ketamine sessions with (*K*-model 7 in the parenthesis, P) or without (*K*-model 4, P and Y/B) additional change in the tendency of choice perseveration (Figure 3, Table 3). Under ketamine, value update increased positively following all outcomes, thus increasing overall probability of repeating the same choice or namely, choice perseveration. However, the change was larger for non-gain outcomes, with the largest modulation after loss, consistent with the results from our model comparison that the model including only a perseveration component performed worse than the best model in all three animals (Figure 3). When the effect of ketamine was simulated with the best models within each class of ketamine models, *K*-model 4 (differential modulation of outcome evaluation for all three outcomes) produced the most similar results to actual data (Figure 4, Supplemental Figure S3).

Unlike monkey Y and B, the best-fitting model for monkey P indicated that ketamine increased overall tendency to switch choice in addition to outcome-dependent modulation of outcome evaluation. However, BIC differed only slightly (dBIC = 3.99) between the best-fitting (*K*-model 7) and the second-best model (*K*-model 4) and the model predictions for choice behavior were very similar both qualitatively and quantitatively (Table 3, Figure 4). We conclude that the behavioral effects of ketamine were consistent across all three monkeys.

We did not find convincing evidence to support that ketamine significantly modulated cognitive (i.e. memory, TCA models) or motivational (i.e. motivation model) aspects of the animal’s behavior. However, ketamine induced ocular nystagmus particularly when animals were holding fixation (i.e. pre-feedback fixation) on a peripheral target immediately after a choice was made.

Centripetal eye drift measured by eye velocity was significantly larger under ketamine (Figure 5; 0.5mg/kg IM, 3-way interaction among drug, dose and time in *ANOVA, F*(3, 523)=20.69, *p*<.001 for P, *F*(3,439)=118.33, *p*<.001 for Y, 2-way interaction between drug and time, *F*(3, 124)=40.72, *p*<.001 for B; 1mg/kg IN, 3-way interaction, *F*(3, 103)=3.54, *p*=.018 for Y, *F*(3,242)=6.26, *p*<.001 for B, main effect of drug, *F*(1, 326)=20.61, *p*<.001 for P). Overall magnitude of nystagmus was significantly larger when ketamine was administered intramuscularly than intranasally (*t*-test, *p*<.001 for all three animals).

**Figure 5.**
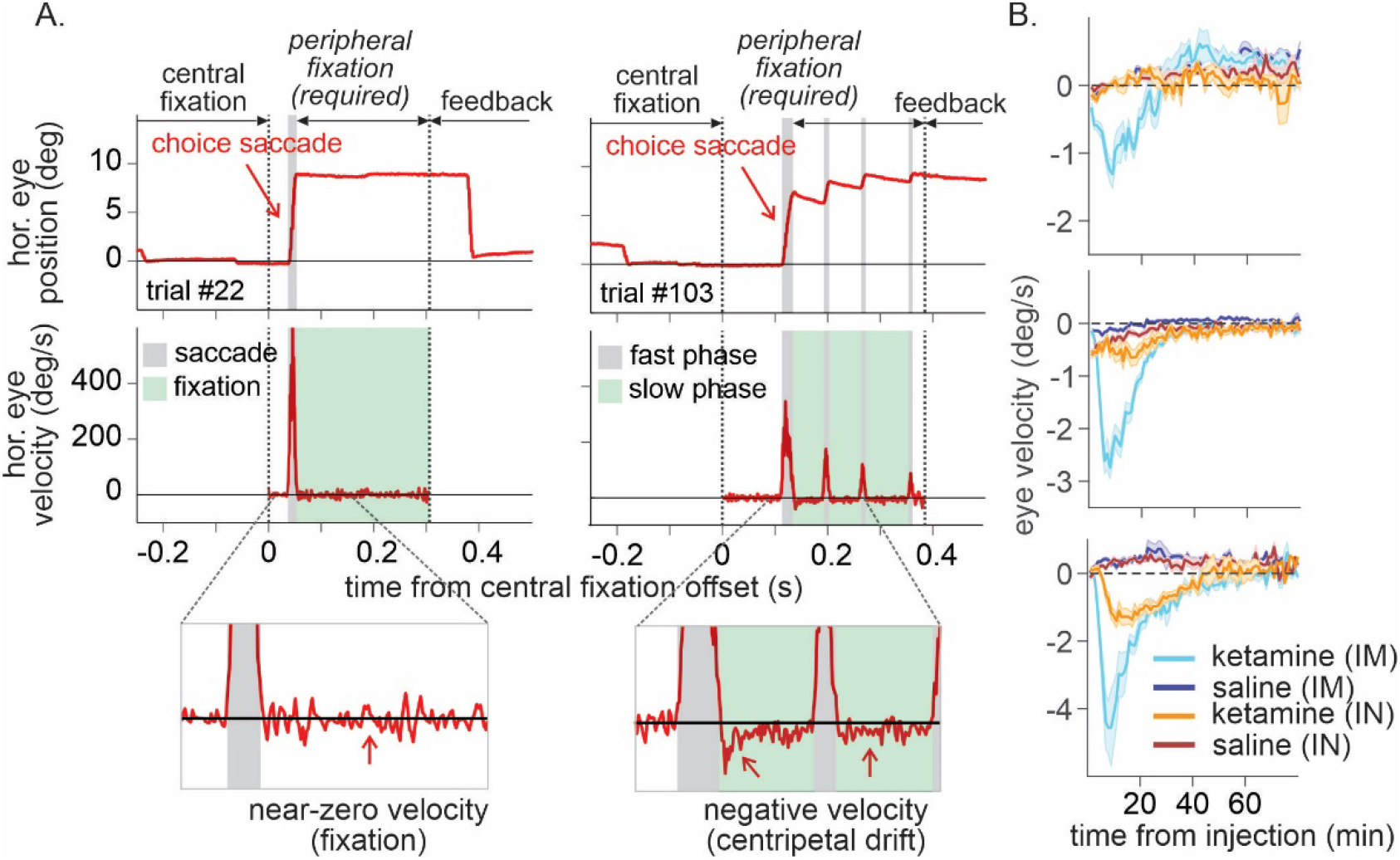
Time course of ketamine-induced ocular nystagmus. **A**. Ocular position and velocity during fixation on the peripheral target in example trials during saline (left panel) and ketamine (right panel) sessions. **B**. Time course of mean ocular velocity aligned at the time of saline or ketamine injection. Shades indicate standard error. IM, IN indicates intramuscular and intranasal administration, respectively.

Ketamine’s effects on both ocular nystagmus and negative evaluation of loss peaked and waned away within 1 hour after the injection (Figure 5, 6). Despite the similar time course, further analysis suggested that these two effects were unlikely to be produced by a common mechanism/process. Nystagmus tended to cause fixation errors during the pre-feedback waiting period in a small number of trials. We found that as the animals owned more tokens and therefore were closer to earning juice reward, fixation errors became less frequent under ketamine (2-way interaction between drug condition and # of owned token in linear regression, *t*=-3.6, -3.3, -9.5 for P, Y and B respectively, *p*<.001 for all three animals; Figure 7, A), suggesting that animals could counter the undesirable effect of ketamine when they had stronger motivation to obtain juice reward. However, ketamine-induced modulation of loss evaluation (i.e. reduced probability of choice switch after loss) did not systematically change with animals’ motivation gated by the number of accrued tokens (2-way interaction between drug condition and # of owned tokens in linear regression, *p*>0.25 for all three animals; Figure 7, B).

**Figure 6.**
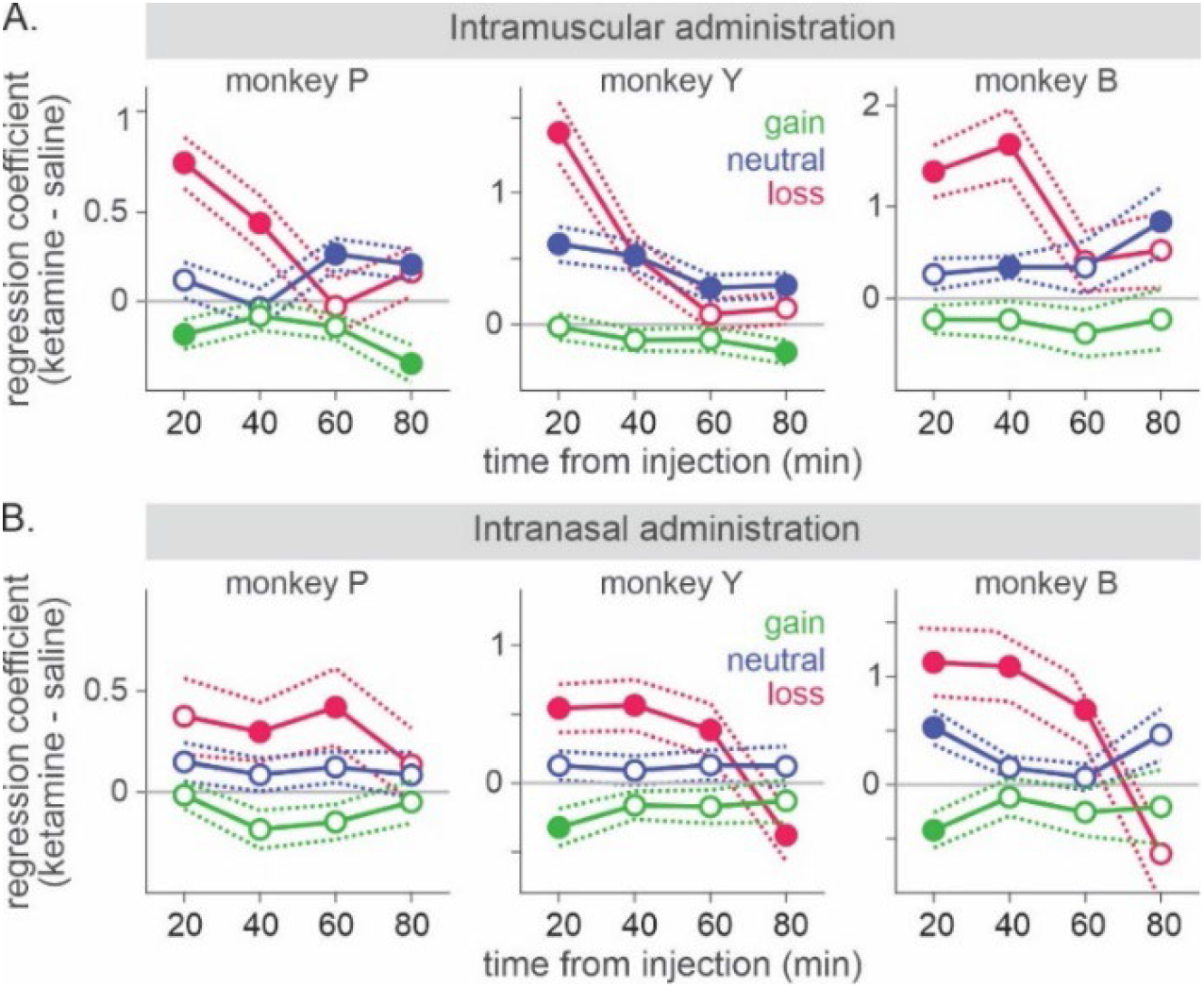
Time course of ketamine-induced modulation of outcome-dependent choice behavior. Time course of attenuation in loss evaluation induced by 0.5mg/kg of intramuscularly (**A**) and 1mg/kg of intranasally (**B**) administered ketamine. Regression coefficient reflecting ketamine’s modulation of the effect of each outcome from the previous trial (trial lag 1) is plotted as a function of time relative to the injection. Dotted lines represent standard error obtained from shuffled data between saline and ketamine sessions separately for each outcome.

**Figure 7.**
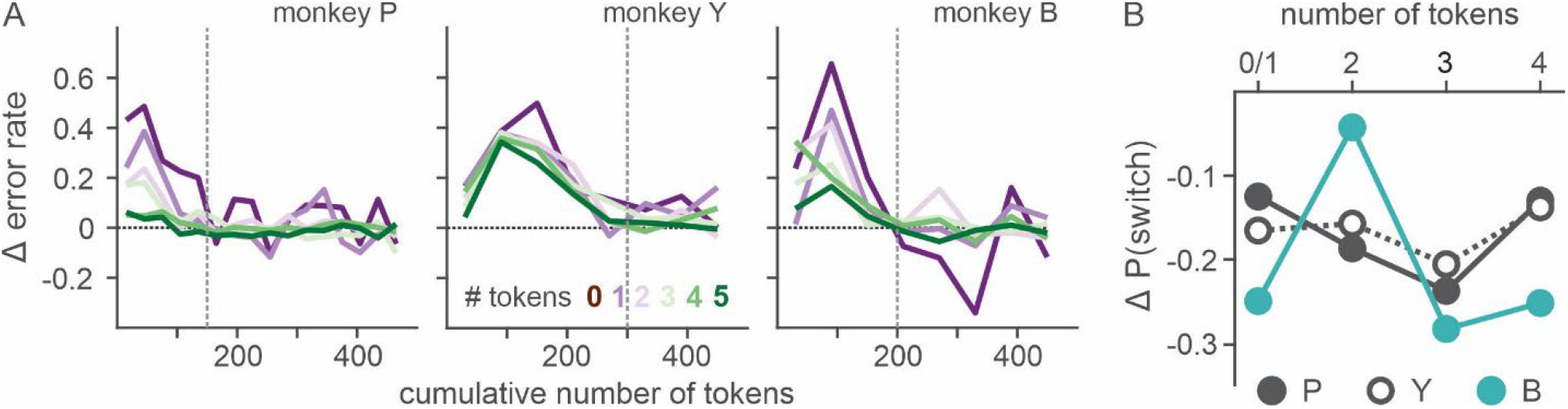
Effect of motivation on countering ketamine-induced fixation errors. **A**. Difference in the rate of fixation break/trial between ketamine and saline sessions is plotted as a function of cumulative number of tokens (as a proxy for time with the satiation effect being controlled). Number of tokens owned by the animal at a given trial (asset) is color-coded. Vertical dotted lines demarcate the latest data point that was included in the linear regression analysis. **B**. Difference in the probability of choice switch after loss from the previous trial (trial lag 1) between ketamine and saline sessions is plotted as a function of asset at the time of decision. Due to limited number of loss trials, analysis was performed after dividing trials into 4 groups according to asset (0∼1, 2, 3, 4).

## Discussion

The acute behavioral effects of ketamine that might reflect antidepressant activity have seldom been systematically examined as the potential antidepressant effects could be obscured by dissociative side-effects in the assessment based on self-report/rating (4, 15, 25, 36). Using a token-based decision task and extensive computational modeling, we examined the behavioral modulation induced by therapeutic doses of ketamine to gain insights into possible early signs of ketamine’s antidepressant activity. We identified the robust effect of ketamine in attenuating negative evaluation of decision outcomes, without affecting the animal’s motivation to obtain juice reward, the ability for proper temporal credit assignment (TCA) or the time constant of memory decay for decision outcomes.

Negative bias in perception, memory, feedback- and emotional processing has been reported to be one of the core ehavioral/neuropsychological features of depression and negative affective bias, in particular, was proposed to play a causal role in the development, maintenance and treatment of depression (37-39). Changes in emotional processing - increased (decreased) positive (negative) affective bias early in the course of antidepressant treatment were also reported to predict later clinical response (40-43). While ketamine’s antidepressant effect is reported to be sustained over a week of period (5), ketamine’s effect on outcome evaluation was acute and did not last over subsequent days (Supplemental Figure S4). This discrepancy might be attributable to the possible differences in the state of brain network between healthy subjects and those with depression as well as the type of measures taken to assess ketamine’s effect. It is noteworthy that even in our task in which the contingency of gains and losses were designed to reinforce dynamic choice, the impact of loss was cumulative as memory decayed slowly over trials and therefore, attenuation of initial impact of loss had a longer-term effect beyond the time of initial evaluation (Figure 1, C). In real-world situations where life events have consequences and their memory decays at much longer time scales than in our study, ketamine’s acute/early effect might be able to mediate longer-term changes in negative affective processing/schemata and mood while patients continue to interact with their social environment during the course of treatment (39, 41, 44). Nevertheless, systematic studies are required to understand whether the reduced aversiveness to loss in our task might share the same mechanisms that underlie ketamine’s antidepressant action.

Ketamine has been used as a pharmacological model of schizophrenia inducing cognitive deficits (14, 45, 46). Ketamine was reported to impair cognitive functions such as working memory, context-dependent behavioral control, task-switching, rule learning and reasoning (47-51). We tested multiple hypotheses regarding ketamine’s possible effects on the time constant of choice and outcome memory, accurate association between an outcome and its causative choice (i.e. temporal credit assignment) and behavioral perseveration. Ketamine increased the overall probability of repeated choice. However, since this tendency was outcome-dependent, with the effect being significantly larger for non-gain outcomes, and largest for loss, this effect cannot be entirely explained by increased perseveration alone. These results are consistent with a previous report that Ketamine didn’t affect the process of learning, but only affected the integration of information for decision (52). These results also suggest that the memory of affective events and their associations (i.e. temporal credit assignment) might be more resistant to acute disruption by NMDA receptor blockade, than affectively neutral information or rules. This disruption-resistant affective memory might underlie the delayed onset of antidepressant’s therapeutic effect and treatment-resistant depression.

It was reported that oculomotor side-effects induced by ketamine rapidly decreased over repeated administrations (53). Interestingly, we found that ketamine-induced fixation errors could be countered at a very short time scale (i.e., on a trial basis) through animals’ strong motivation to earn juice reward, further corroborating our findings that ketamine didn’t change the animal’s motivation for reward.

Finally, our results demonstrate the sensitivity and the potential of the non-human primate model for investigation of the neurobiology of depression and antidepressant treatments, warranting further study on the mechanisms of ketamine’s antidepressant action over different mood states and contexts carrying affective events that have impacts at diverse time scales.

## Acknowledgements

We thank Melissa Boucher and Khalil AbedRabbo for their technical assistance. We also thank Drs. Daeyeol Lee and Amy Arnsten for their helpful discussion and thoughtful comments throughout the study. This work was supported by the National Institute of Health (R01 MH108643).

## Supplemental Information

### Analysis of plasma concentration of ketamine

To be able to effectively compare behavioral effects observed following intramuscular (IM) and intranasal (IN) administration of ketamine, we measured plasma concentration of ketamine following IM- and IN-administrations in monkey B (1 session for 0.5mg/kg IM; 2 sessions for 0.5mg/kg IN; 2 sessions for 1mg/kg IN administration). Blood samples were collected at 10, 25, 40, 60 and 75 minutes following ketamine administration and analyzed with the HPLC protocol modified from Gross et al., (1999) (1). 100ng of the internal standard clonidine and 0.5ml of thawed plasma were added to a 15ml polypropylene tube and 15ml of diethyl ether added. Following 5 min of vortex mixing, 4 mL of the upper ether layer was transferred to a new polypropylene tube and 250 µL of 0.01 M phosphoric acid added. After vortex mixing for 5 min, the lower aqueous layer was transferred to a 1.5 mL tube and evaporated under vacuum (∼ 1 hour, SpeedVac evaporator, medium heat). The dry residue was dissolved in 150 μL of 0.01 sodium monobasic dihydrogen phosphate. 50 μL of the extract was injected and separated on a 3 μm Cyano Spherisorb HPLC column (4.6 x 100 mm) eluted with 80% 0.01 sodium acetate/20% acetonitrile (0.8 mL/min, 40C). Norketamine, ketamine and clonidine were detected by UV absorption (220 nm; Hitachi L-7400) with retention times of 6.1, 8.1 and 11.0 minutes, respectively; and with absolute detection limits of 0.2, 0.5 and 0.3 ng, respectively. Extracted standards were prepared using 0.5 mL aliquots of pooled human plasma. Plasma drug concentrations of samples were calculated knowing the peak height ratios of the extracted standards, the peak height ratios in samples and the amount of internal standard added (200 ng/mL). When injecting 50 μL of the extract, norketamine and ketamine were determined with concentration detection limits of 1 and 2 ng/mL, respectively.

**Supplemental Figure S1.**
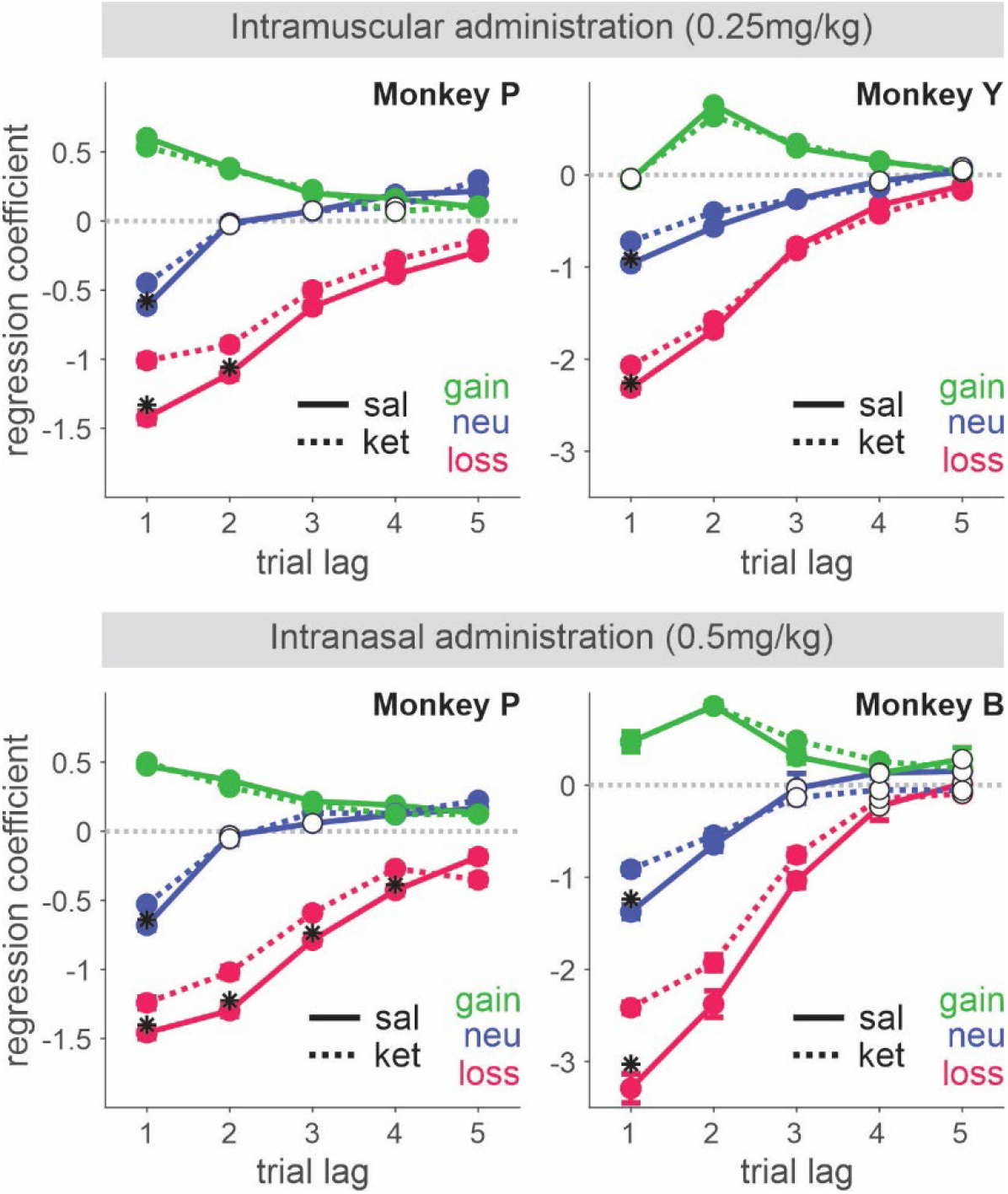
Behavioral effect of 0.25mg/kg intramuscularly (IM) administered (top) and 0.5mg/kg intranasally (IN) administered (bottom) ketamine. Regression coefficients reflecting the effects of gain, neutral (zero-token) and loss outcomes obtained in the past trials are plotted. Solid and dotted lines represent data from saline and ketamine sessions respectively. Solid (empty) symbols indicate that the corresponding coefficients are (not) significantly different from zero. * indicates that the coefficients from ketamine and saline sessions are significantly different for corresponding trial lag of outcome.

**Supplemental Figure S2.**
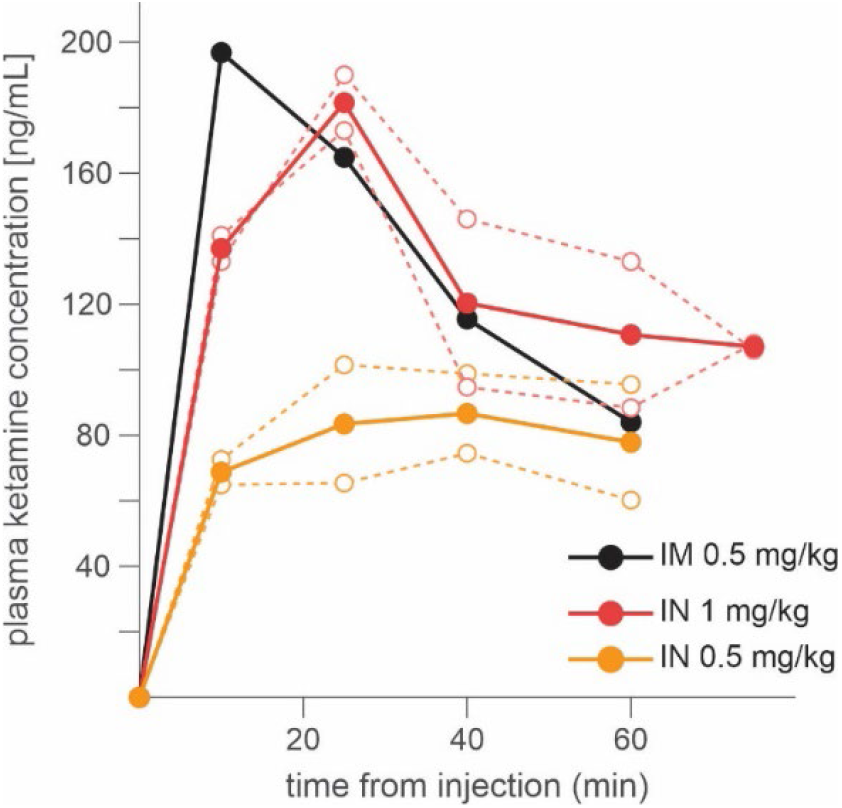
Time course of plasma concentration of ketamine following intramuscular (IM) and intranasal (IN) administration. Dotted lines indicate the data from individual sessions, and dotted lines for average across individual sessions. Blood sample was taken every 20 minute after injection.

**Supplemental Figure S3.**
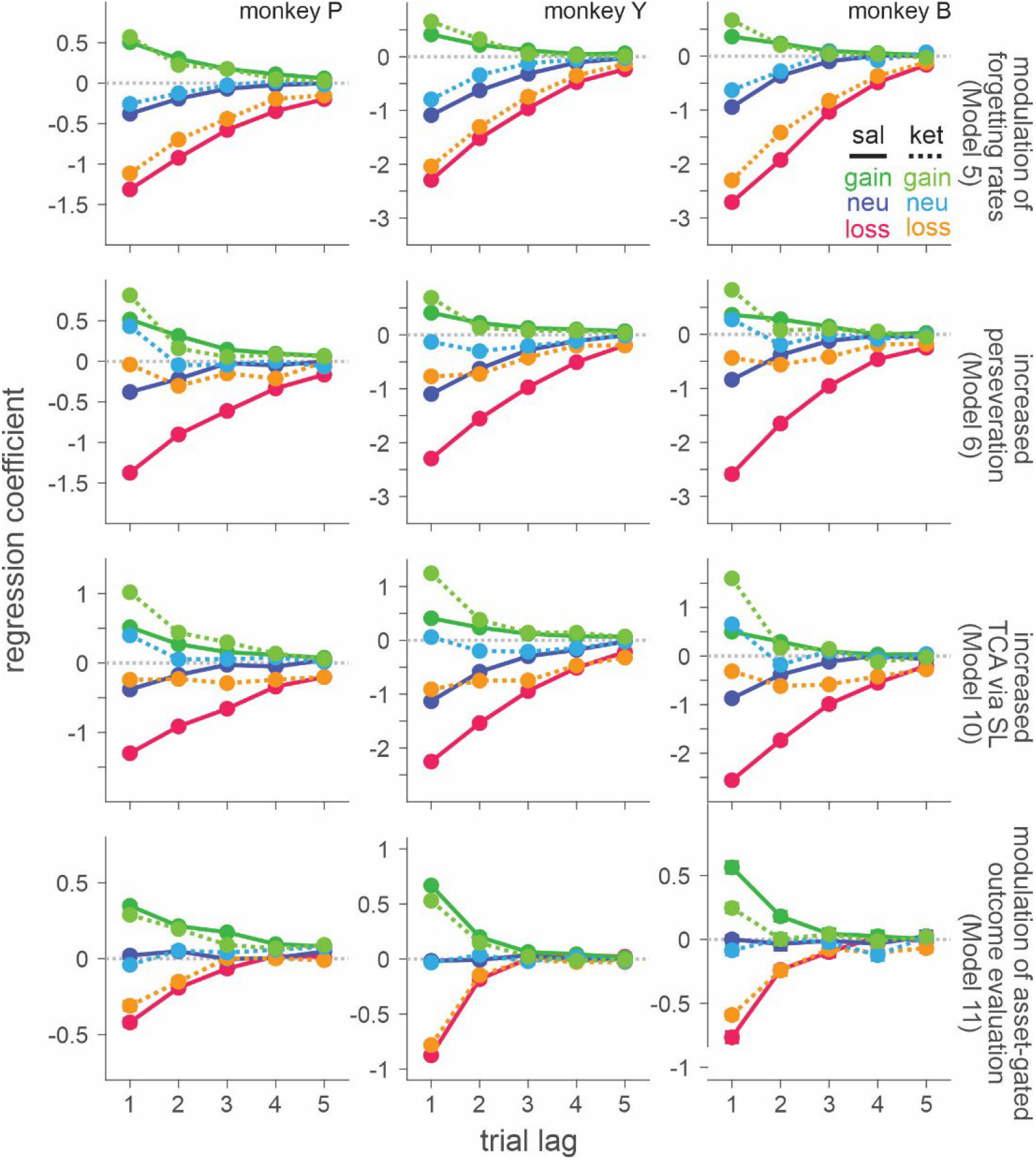
ketamine-induced behavioral modulation simulated with best-fitting *K-*model of each class of reinforcement learning (RL) models. Format is same as in Figure 4. Each column represents simulated data with best-fitting parameters for individual monkey (P, Y and B from left to right).

**Supplemental Figure S4.**
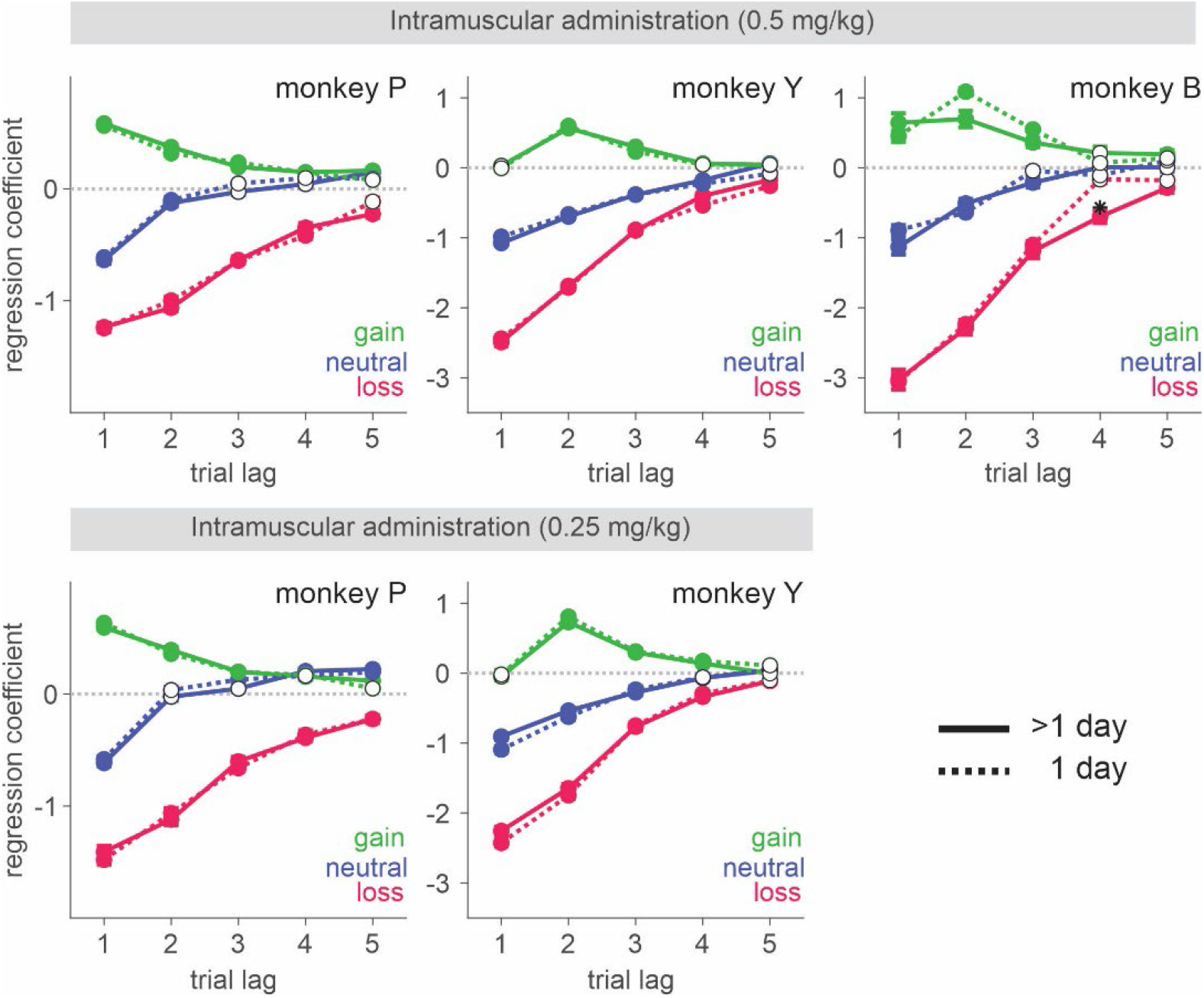
Behavioral effects of ketamine do not spread over the sessions subsequent to the injection. Regression coefficients reflecting the effects of gain, neutral (zero-token) and loss outcomes obtained in the past trials are plotted. Solid (dotted) lines represent data from saline sessions > 1 day (1 day) after a ketamine session. Solid (empty) symbols indicate that the corresponding coefficients are (not) significantly different from zero.

